# The Glycolytic Metabolite Methylglyoxal Covalently Inactivates the NLRP3 Inflammasome

**DOI:** 10.1101/2024.04.19.589802

**Authors:** Caroline Stanton, Chavin Buasakdi, Jie Sun, Ian Levitan, Prerona Bora, Sergei Kutseikin, R. Luke Wiseman, Michael J. Bollong

## Abstract

The NLRP3 inflammasome promotes inflammation in disease, yet the full repertoire of mechanisms regulating its activity are not well delineated. Among established regulatory mechanisms, covalent modification of NLRP3 has emerged as a common route for pharmacological inactivation of this protein. Here, we show that inhibition of the glycolytic enzyme PGK1 results in the accumulation of methylglyoxal, a reactive metabolite whose increased levels decrease NLRP3 assembly and inflammatory signaling in cells. We find that methylglyoxal inactivates NLRP3 via a non-enzymatic, covalent crosslinking-based mechanism, promoting inter- and intra-protein MICA posttranslational linkages within NLRP3. This work establishes NLRP3 as capable of sensing a host of electrophilic chemicals, both exogenous small molecules and endogenous reactive metabolites, and suggests a mechanism by which glycolytic flux can moderate the activation status of a central inflammatory signaling pathway.

## Introduction

Vital to regulating the levels of pro-inflammatory cytokines and ultimately pyroptotic cell death, inflammasomes are large cytosolic protein complexes controlled by pattern recognition receptors. One such inflammasome is nucleated by NLRP3 (NOD-, LRR- and pyrin domain-containing protein 3), which responds to various danger and pathogen associated molecular patterns. Activation of the NLRP3 inflammasome requires both priming, activation of the transcription factor NF-κB to augment transcription of inflammasome components, as well as a secondary activating signal, such as K^+^ efflux, lysosomal signaling, or mitochondrial reactive oxygen species. The presence of both signals promotes assembly of the active NLRP3 inflammasome, a complex formed by association of NLRP3 with NEK7, ASC, and pro-caspase 1. Upon assembly, pro-caspase-1 activation cleaves pro-IL-1β and pro-IL-18 to active forms. Additionally, caspase 1 cleaves gasdermin D which forms pores in the plasma membrane resulting in pyroptotic cell death. Given the broad array of signals to which NLRP3 can respond, the NLRP3 inflammasome is frequently associated with sterile inflammation in disease^1-7^. For this reason, NLRP3 remains a promising drug target with the potential to mitigate pathology in numerous inflammatory conditions. However, at present, there are no clinically approved NLRP3 inhibitors, though several are being evaluated for efficacy in the clinic^8,9^.

Dapansutrile and ZYIL1, inhibitors targeting the NACHT domain of NLRP3 which reduces ATPase activity, have shown promising activity in phase 2 studies ^9-11^. However, among promising approaches to inhibit NLRP3 pharmacologically, covalent inhibitors of NLRP3 have become commonplace in the clinic and in literature. RRx-001, a covalent modifier of C409 which inhibits protein interactions with NEK7, is in phase 3 studies for cancer, but since its identification as an inflammasome inhibitor, is now also being studied for various neurodegenerative diseases^12^. Likewise, other NLRP3 inhibitors targeting several reactive sensor cysteines in NLRP3 have been reported in recent years, including oridonin, costunolide, Bay-11-7082, MNS, parthenolide, and Tubocapsanolide A^13-17^. Recently, we performed an unbiased high-throughput screen which resulted in the identification of nine new covalent scaffolds with three of these chemical scaffolds showing robust covalent modification of NLRP3 across several of its domains^18^. Previously NLRP3 has been shown to be inhibited through covalent modification by several reactive metabolites including diroximel fumarate^18^, a fumarate derivative, as well as itaconate at which modifies at C548 of murine NLRP3 which inhibits NLRP3 interactions with NEK7^19^. Collectively, these observations provide mounting evidence that NLRP3 serves as an electrophile sensor within the cell, linking the covalent modification of its sensor cysteines to decreased NLRP3 inflammasome activation and downstream inflammatory signaling.

The capacity for NLRP3 to respond to electrophilic small molecules is reminiscent of the established electrophile sensor KEAP1 (Kelch-like ECH-associated protein 1). KEAP1 serves as a negative regulator of the oxidative stress responsive transcription factor NRF2 (NFE2L2, NFE2 Like BZIP Transcription Factor 2), promoting its ubiquitination and degradation through interactions with the Cullin-RING E3 ubiquitin ligase protein complex. KEAP1 bears 12 sensor cysteines which respond to covalent modification by liberating NRF2 sufficiently to enact a protective transcriptional response^20^. In addition to exogenous electrophilic chemicals, KEAP1 has been shown to covalently sense endogenous reactive electrophilic metabolites induced by perturbations in the TCA cycle and glycolysis, resulting in NRF2 activation^21,22^. Both the TCA cycle metabolite fumarate and the TCA cycle derived metabolite itaconate are established activators of NRF2 that covalently modify sensor cysteines in the BTB domain of KEAP1^23-27^.

We previously identified the compound CBR-470-1 as an activator of NRF2^28^. This compound inhibits the glycolytic enzyme phosphoglycerate kinase (PGK1), leading to an accumulation of upstream metabolites 1,3-bisphosphoglycerate (1,3-BPG), glyceraldehyde-3-phosphate (GAP), and dihydroxyacetone phosphate (DHAP). In response to PGK1 inhibition, methylglyoxal (MGO), the non-enzymatic dicarbonyl elimination product of GAP and DHAP, accumulates and induces formation of KEAP1 dimers via MICA (methyl imidazole crosslink between cysteine and arginine) modifications, augmenting NRF2 activation and expression of its target genes. An analog of this compound, CBR-470-2, was also shown to be active in vivo, protecting against oxidative damage in response to pathogenic UV exposure^28^.

Because NLRP3 and KEAP1 both sense the presence of electrophilic chemicals from exogenous and endogenous sources, we hypothesized that NLRP3 might also be capable of sensing and responding to glycolytic metabolites, like MGO. Here we report that the CBR-470 series of compounds inhibit NLRP3 inflammasome assembly through a mechanism that involves covalent crosslinking and inactivation of NLRP3 by increased levels of MGO.

## Results

### Pharmacologic PGK1 inhibitors block NLRP3 inflammasome assembly and activity

The assembly and activity of the NLRP3 inflammasome is highly sensitive to reactive metabolites generated by alterations to metabolic pathways. Here, we sought to determine whether PGK1 inhibitors such as CBR-470-1 and CBR-470-2 (**Fig. S1A,B**) – both compounds that lead to generation of the reactive metabolite MGO^28^ – could inhibit NLRP3 inflammasome assembly and activity. Initially, we monitored assembly of the NLRP3 inflammasomes in THP1 cells stably expressing a GFP-tagged ASC (ASC-GFP) (Invivogen). Treatment of these cells with the priming stimulus LPS followed by the activating stimulus nigericin triggers NLRP3 inflammasome assembly, which can be followed by the formation of ASC-GFP specks using high content imaging (**Fig. S1C**). As described previously, ASC-GFP speck formation could be dose-dependently inhibited by pre-treatment with the NLRP3 inflammasome inhibitor MCC950 (**Fig. 1A**)^18^. Intriguingly, pretreatment with either CBR-470-1 or CBR-470-2 for 2 h inhibited ASC-GFP speck formation in these cells with an IC50 of 25 µM and 3 µM, respectively (**Fig. 1A**). The levels of inhibition afforded by these two PGK1 inhibitors was similar to that observed for MCC950 and minimal increases in inhibition were observed by increasing the pre-treatment time beyond 2 h (**Fig. 1A-C, Fig. S1D, E**).

**Figure 1.**
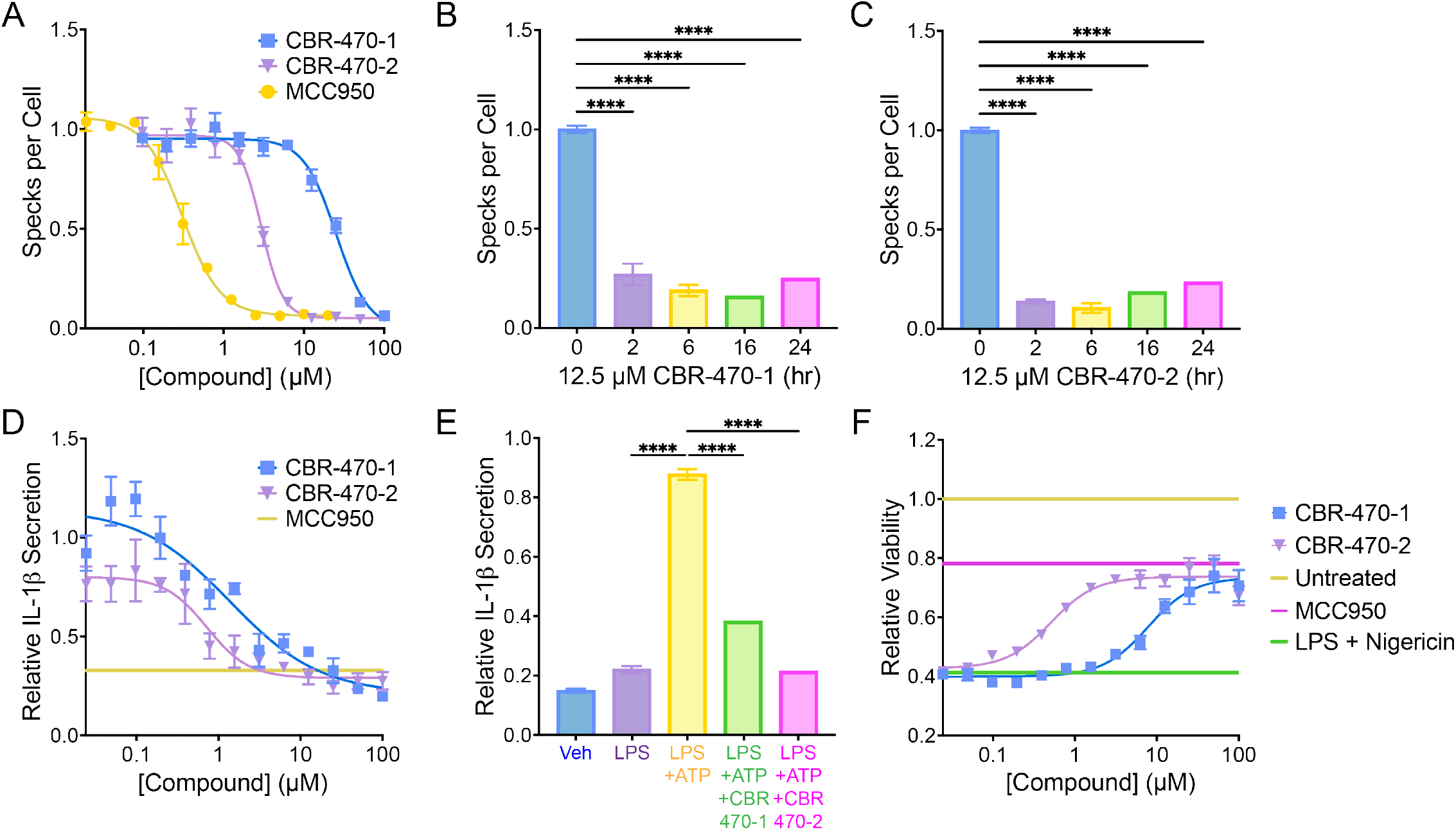
CBR-470 series compounds inhibit the NLRP3 inflammasome. (A) Number of ASC-GFP specks per cell in THP1-ASC-GFP cells treated for 2 hours with the indicated concentrations of compound. Error bars show SEM for n = 3 replicates. (B, C) Number of ASC-GFP specks per cell in THP1-ASC-GFP cells pretreated with CBR-470-1 (B) and CBR-470-2 (C) at 12.5 µM for 0-24 h. Error bars show SEM for n = 3 replicates. \*\*\*\**P*<0.0001 for ordinary one-way analysis of variance (ANOVA). (D) Relative level of secreted IL-1β from WT THP1 cells stimulated with LPS, pretreated with MCC950 (10 µM), CBR-470-1, or CBR-470-2, and activated with ATP. IL-1β measured by SEAP secretion from HEK-Blue-IL-1β reporter cells. Error bars show SEM for n = 3 replicates. (E) Relative level of secreted IL-1β from primary murine dendritic cells stimulated with LPS and pretreated with 10 µM CBR-470-1 or CBR-470-2 for 1 hr prior to addition of ATP as measured by SEAP secretion from HEK-Blue-IL-1β reporter cells. Error bars show SD for n = 16 replicates. \*\*\*\**P*<0.0001 for ordinary one-way analysis of variance (ANOVA) with Tukey correction for multiple comparisons between conditions. (F) Relative viability of LPS-primed (1 µg/mL, 16 h) WT THP1 following NLRP3-mediated pyroptotic cell death induced by Nigericin (10 µM, 2.5 h), pretreated with CBR-470-1 and CBR-470-2 (2 h) in dose response, or with 10 µM MCC950 (2 h). Error bars show SEM for n = 6 replicates.

Next, we determined the potential for these compounds to inhibit downstream NLRP3 inflammasome-dependent activities including IL-1β secretion and pyroptotic cell death. Pretreatment with CBR-470-1 or CBR-470-2 for 2 h inhibited IL-1β secretion from both THP1 cells and primary dendritic cells, as measured using HEK293Blue IL-1β reporter cells (**Fig. 1D,E**). Similarly, pretreatment with CBR-470-1 or CBR-470-2 for 2 h blocked pyroptotic cell death induced by nigericin in LPS-primed THP1 cells to the same levels observed for MCC950 (**Fig. 1F**). These results demonstrate that two different PGK1 inhibitors block NLRP3 inflammasome assembly and downstream signaling. Since CBR-470-2 showed superior potency in cell-based experiments as compared to CBR-470-1, we specifically focused on CBR-470-2 for further mechanistic experimentation.

### CBR-470-2-dependent NLRP3 inhibition is independent of NRF2 Activation or NF-κB Inhibition

PGK1 inhibitors such as CBR-470-2 have been previously reported to activate NRF2 in mammalian cells^28^. Since NRF2 activation has previously been shown to inhibit the NLRP3 inflammasome^29,30^, we sought to determine whether the inhibition of NLRP3 assembly and activity observed with this compound could be attributed to NRF2 activation. Interestingly, a 2 h treatment with CBR-470-2 did not induce expression of the NRF2 target gene *NQO1* in THP1 cells (**Fig. 2A**). However, we did observe increased *NQO1* expression following 16 h treatment with CBR-470-2. This increase in *NQO1* expression could be blocked by co-treatment with the NRF2 inhibitor ML385, indicating that this effect can be attributed to NRF2 activation. Similar results were observed for the alternative NRF2 target gene *HMOX1* (**Fig. S2**). These results indicate that NRF2 is not activated in the 2 h pretreatment time course where we observe CBR-470-2-dependent inhibition of NLRP3 inflammasome assembly and activation.

**Figure 2.**
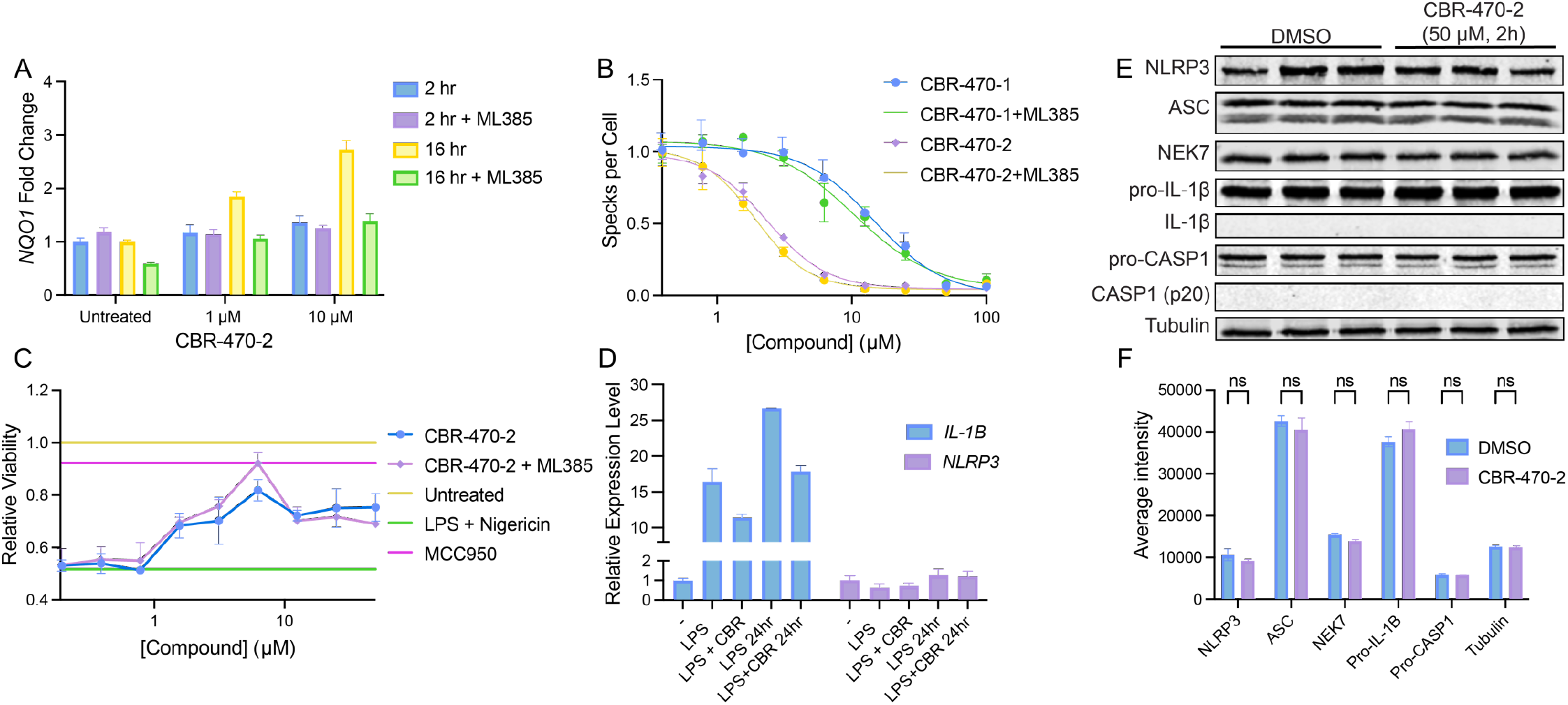
CBR-470-2 inhibition of the NLRP3 inflammasome does not rely on NRF2 activation or NF-κB inhibition. (A) Relative transcript level for *NQO1* as measured by qPCR from WT THP1 cells pre-treated with or without 10 µM ML385 for 30 min and then treated CBR-470-2 for 2 or 16 h. Error bars show SEM for n = 3 replicates. (B) Number of ASC-GFP specks per cell in THP1-ASC-GFP cells pretreated with or without 10 µM ML385 for 30 min and then for 2 hours with CBR-470-1 and CBR-470-2. Error bars show SEM for n = 3 replicates. (C) Relative viability of LPS-primed (1 µg/mL, 16 h) WT THP1 following NLRP3-mediated pyroptotic cell death induced by Nigericin (10 µM, 2.5 h), pretreated with or without 10 µM ML385 for 30 min and then with CBR-470-2 (2 h) in dose response, or with 10 µM MCC950 (2 h). Error bars show SEM for n = 3 replicates. (D) Relative transcript levels for NF-κB target genes *NLRP3* and *IL-1β* as measured by qPCR from WT THP1 cells treated with 1 µg/mL LPS for 3 hr and then 10 µM CBR-470-2 for 3 or 24 h. Error bars show SEM for n = 3 replicates. (E) Western blot for NLRP3, ASC, NEK7, pro-IL-1β, IL-1β, pro-CASP1, CASP1, and Tubulin in LPS-primed THP1 cells treated with vehicle or CBR-470-2 (50 µM) for 2 h. (F) Average intensity of bands of western blot in E for each protein.

To further probe the specific dependence of compound-dependent NLRP3 inhibition, we monitored ASC-GFP speck formation and pyroptotic cell death in THP1 cells pretreated with the NRF2 inhibitor ML385 for 30 min followed by CBR-470-2 for 2 h. Interestingly, pre-treatment with ML385 did not influence the potency or efficacy of CBR-470-2-dependent inhibition of NLRP3 inflammasome assembly or activity in these assays (**Fig. 2B,C**). This further demonstrates that the observed inhibition of NLRP3 inflammasomes afforded by the PGK1 inhibitor is not mediated through NRF2 activation.

Next, we sought to determine the potential impact of CBR-470-2 on the NF-κB-dependent expression of NLRP3 inflammasome components induced by LPS priming. We found that co-treatment with CBR-470-2 for 3 h or 24 h does not significantly influence expression of *NLRP3* in LPS-primed THP1 cells, although it does modestly reduce expression of *pro-IL1b* (**Fig. 2D**). Further, we found protein levels of NF-κB targets including NLRP3, ASC, and pro-IL-1β, as well as other NLRP3 inflammasome components were not significantly decreased following a 2 h treatment with CBR-470-2 in LPS-primed THP1 cells (**Fig. 2E, F**). These results indicate that CBR-470-2 does not significantly influence expression of NF-κB target genes induced by LPS priming.

### CBR-470-2 inhibits NLRP3 through the increased production of MGO downstream of PGK1

Pharmacologic PGK1 inhibition with CBR-470-2 leads to the accumulation of the reactive metabolite MGO^28^. Since previous results showed that the NLRP3 inflammasome is sensitive to inactivation by reactive metabolites, we predicted that CBR-470-2 likely inhibits NLRP3 inflammasome assembly and activity through a mechanism involving the accumulation of MGO. To initially test this, we monitored NLRP3 inflammasome assembly (via ASC-GFP speck formation) in THP1 cells co-treated with CBR-470-2 and glutathione (GSH), the latter a treatment that neutralizes reactive metabolites. Intriguingly, we found that co-treatment with glutathione fully abrogated the inhibition of ASC-GFP speck formation afforded by CBR-470-2 (**Fig. 3A**). Identical results were observed in THP1 cells treated with CBR-470-1 (**Fig. S3A**). This supports a model whereby these two PGK1 inhibitors inhibit inflammasome assembly through a mechanism involving the accumulation of a reactive metabolite such as MGO.

**Figure 3.**
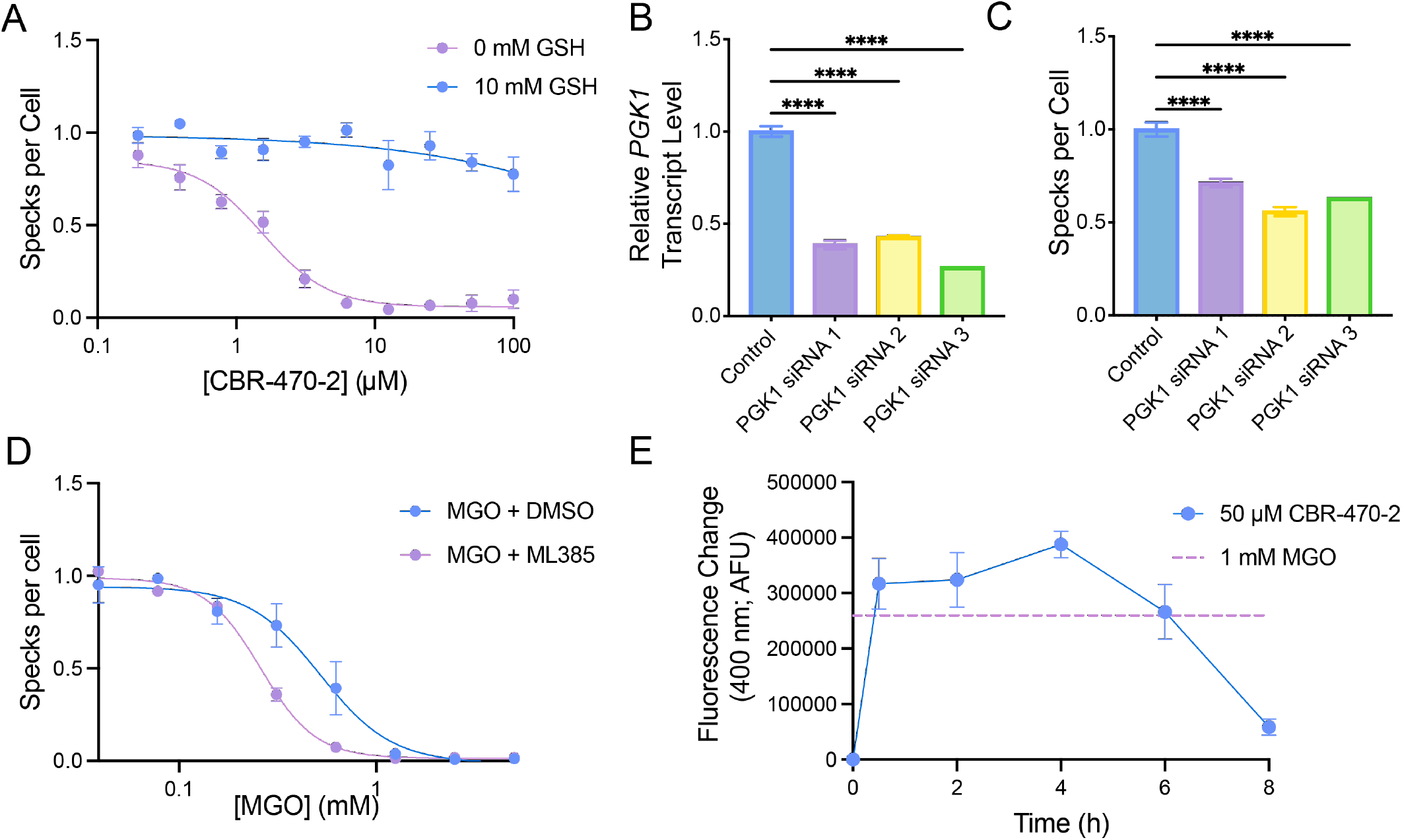
Inhibition by CBR-470 is mediated by PGK1 inhibition and accumulation of methylglyoxal. (A) Number of ASC-GFP specks per cell in THP1-ASC-GFP cells pre-treated with 0 or 10 mM GSH and then treated in dose response with CBR-470-2. Error bars show SEM for n = 3 replicates. (B) Relative transcript level of *PGK1* as measured by qPCR 48 hr after knockdown with siRNA targeting *PGK1* in THP1-ASC-GFP cells. Error bars show SEM for n = 3 replicates (C) ASC Speck formation in THP1-ASC-GFP cells 48 hr following transfection with siRNAs targeting *PGK1*. Error bars show SEM for n = 3 replicates. (D) Number of ASC-GFP specks per cell in THP1-ASC-GFP cells pretreated with or without 10 µM ML385 for 30 min and then for 2 hours with MGO in dose response. Error bars show SEM for n = 3 replicates. (E) Fluorescence (Ex. 355, Em. 400 nM) increase from untreated baseline control for methylglyoxal level assay for lysates from THP1 cells treated with 50 µM CBR-470-2 for indicated time point or 1 mM MGO for 1 h. Error bars show SEM for n = 6 replicates.

CBR-470-2 generates MGO by inhibiting the enzymatic activity of PGK1^28^. Thus, we predicted that genetic depletion of PGK1 should similarly inhibit NLRP3 inflammasome assembly. We depleted *PGK1* in THP1-ASC-GFP cells using three different siRNAs and monitored NLRP3 inflammasome assembly by ASC-GFP speck formation. We confirmed that our siRNA treatment reduced expression of *PGK1* in these cells by RT-qPCR (**Fig. 3B**). Intriguingly, reductions in *PGK1* afforded by all three siRNAs significantly reduced ASC-GFP speck formation in THP1 cells (**Fig. 3C**). This indicates that genetic depletion of *PGK1* recapitulates the reduced NLRP3 inflammasome assembly induced by pharmacologic PGK1 inhibitors such as CBR-470-2.

Next, we determined the potential for direct administration of MGO to inhibit NLRP3 inflammasome. We treated THP1 cells with increasing concentrations of MGO and monitored NLRP3 assembly using our ASC-GFP speck formation assay. Pre-treatment with MGO for 2 h inhibited NLRP3 inflammasome assembly with an EC50 of ∼0.52 mM (**Fig. 3D**). Similar results were observed for pyroptotic cell death, where pretreatment for 2 h with MGO similarly improved viability of THP1 cells treated with LPS and nigericin (**Fig. S3B**). MGO activates NRF2 through the covalent targeting of KEAP1. Thus, we determined the potential involvement of MGO-dependent NRF2 activation in the inhibition of NLRP3 inflammasome assembly afforded by this metabolite by pretreating THP1 cells with the NRF2 inhibitor ML385 prior to MGO. We found that pre-treatment with ML385 caused a modest increase in dose-dependent inhibition of ASC-GFP speck formation afforded by MGO treatment (**Fig. 3D**). Similarly, co-treatment with ML385 also caused a modest increase in the dose-dependent protection against pyroptotic cell death induced by MGO (**Fig. S3B**). These results suggests that MGO-dependent NRF2 activation does not contribute to the observed reduction in NLRP3 inflammasome assembly and activity induced by this metabolite, and our results are also consistent with a model whereby MGO inhibits NLRP3 inflammasome assembly through a direct mechanism involving covalent protein modification.

We next sought to determine whether treatment with CBR-470-2 increases MGO levels in THP1 cells. Towards that aim, we used a fluorescence indicator probe for MGO generation^31^ to monitor the production of this metabolite in THP1 cells treated with CBR-470-2 (**Fig. S3C**). We observed a rapid increase of MGO in THP1 cells treated with CBR-470-2, which began to decline ∼8 h after treatment (**Fig. 3E**). Intriguingly, the level of intracellular MGO generated by compound treatment is nearly identical to that observed in THP1 cells treated directly with 1 mM MGO – a dose sufficient to fully inhibit NLRP3 inflammasome assembly (**Fig. 3D**). We also observe crosslinking of KEAP1 following treatment of THP1 cells with CBR-470-2 within 1 h, supporting this rapid accumulation of MGO within the cell (**Fig. S3D**). This indicates that treatment with CBR-470-2 increases intracellular MGO to levels sufficient to inhibit NLRP3 inflammasome assembly.

Collectively, these results indicate that CBR-470-2 inhibits NLRP3 inflammasome assembly through a mechanism involving PGK1 inhibition and subsequent accumulation of the reactive metabolite MGO.

### MGO induces inter- and intra-molecular crosslinks of NLRP3 monomers through MICA modifications

MGO inhibits the NRF2 suppressor KEAP1 by inducing MICA (methyl imidazole crosslink between cysteine and arginine) modifications on neighboring protomers. Thus, we sought to determine whether MGO generated by CBR-470-2 treatment could similarly induce intermolecular crosslinking of NLRP3. Treatment with MGO for 2 h increased crosslinking of NLRP3-FLAG expressed in HEK293T cells, as measured by immunoblotting, evident by the accumulation of high molecular weight (HMW) NLRP3 (**Fig. S4A**). Similarly, treatment with CBR-470-2 also increased HMW NLRP3-FLAG in HEK293T cells. Further, treatment with CBR-470-2 increased HMW endogenous NLRP3 in THP1 cells, evident by reductions in soluble NLRP3 and increased HMW NLRP3 in the insoluble fraction (**Fig. 4A**).

**Figure 4.**
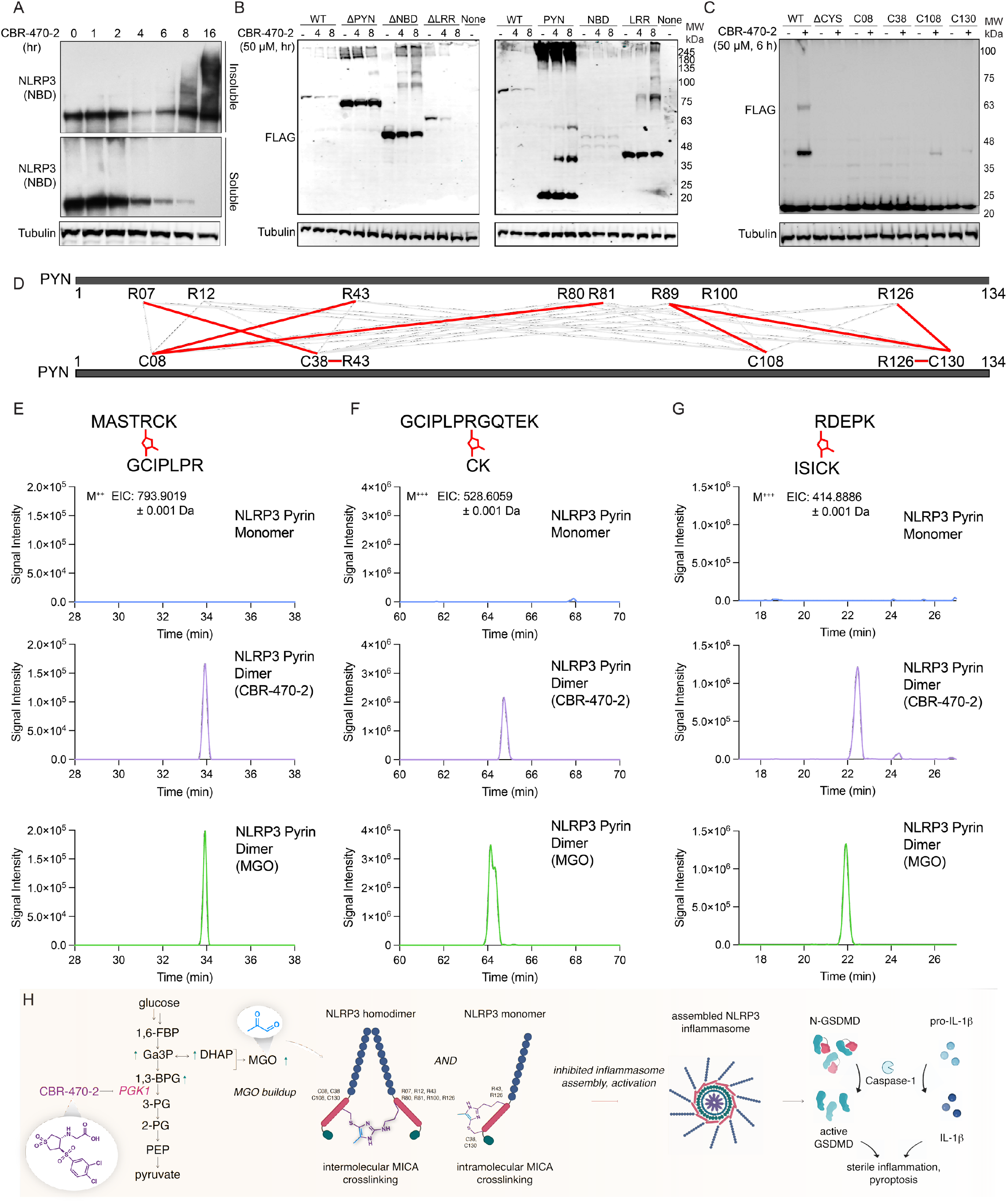
CBR-470-2 inhibits NLRP3 through MICA crosslinks. (A) Western blot of Soluble and Insoluble NLRP3 (NBD) and Tubulin from WT THP1 cells treated with 50 µM CBR-470-2 for 1 to 16 h. (B) Western blot for FLAG and Tubulin in HEK293T cells overexpressing the indicated FLAG-Tagged NLRP3 domain constructs treated with 50 µM CBR-470-2 for 4 or 8 h. (C) Western blot of FLAG from HEK293T cells overexpressing NLRP3-FLAG PYN domain constructs with cysteines mutated and individually reintroduced, treated with 50 µM CBR-470-2 for 6 h. (D) Schematic depicting the various cysteine-arginine MICA crosslinks observed within NLRP3 PYN domain (E-G) EICs from LC–MS/MS analyses of gel-isolated and digested HMW-PYN (CBR-470-2 and MGO-induced) and monomeric PYN for C38–R07 (E), C08-R43 (F), and C130-R89 crosslinked peptides. (H) Schematic depicting the communication between glucose metabolism and the NLRP3 inflammasome mediated by MGO crosslinking of NLRP3 and inhibition of inflammasome assembly.

We next sought to identify the specific domains of NLRP3 that were most sensitive to crosslinking induced by treatment with CBR-470-2 or exogenous MGO. NLRP3 contains multiple domains including a pyrin domain (PYN), a nucleotide binding domain (NBD), and a leucine rich repeat (LRR) domain. We expressed NLRP3-FLAG constructs lacking each of these individual domains (e.g., βPYN, βNBD, βLRR) or each individual domains on their own (e.g., PYN, NBD, LRR) in HEK293T cells and monitored crosslinking induced by treatment with CBR-470-2 or MGO. Interestingly, all of these constructs demonstrated increases in HMW bands in cells treated with either CBR-470-2 or MGO (**Fig. 4B, Fig. S4B**). However, loss of the PYN domain from the full-length NLRP3-FLAG construct appeared to most strongly inhibit crosslinking under both conditions. Likewise, when expressed in isolation, the PYN domain showed the most robust increase in HMW bands when compared to other domains expressed on their own. Intriguingly, we found that recombinant PYN (residues 3-110) also readily form HMW bands corresponding to molecular weights of PYN dimers and trimers when treated with 1 mM MGO for 1 h in vitro (**Fig. S4C-E**). Thus, we explicitly sought to probe the nature of this modification using the PYN domain construct.

MICA modifications proceed by crosslinking Cys residues to Arg residues. Thus, we overexpressed PYN domains lacking all or 3 of the 4 Cys residues within this domain in HEK293T cells and monitored crosslinking induced by treatment with CBR-470-2 by immunoblotting. This showed that loss of all Cys residues prevented CBR-470-2-dependent increases in PYN crosslinking (**Fig. 4C**). Intriguingly, reintroduction of C108 or C130 into the Cys-less PYN domain partially restored CBR-470-2-dependent crosslinking. Similar results were observed in MGO cells overexpressing these PYN Cys mutants, although all constructs showed some modest increase in MGO-dependent crosslinking (**Fig. S4F**). These results identify C108 and C130 of the PYN domain as two sites for covalent modification. Using two alkyne containing probe compounds, mCMF859 and P207-9175, which previously were shown to inhibit NLRP3 and covalently modify C130 in the PYN domain (**Fig. S4G**), we further confirmed that CBR-470-2 treatment decreases probe labeling of this Cys residue (**Fig. S4H**). This further supports a model whereby MGO generated by CBR-470-2-dependent PGK1 inhibition leads to the covalent modification of this site.

Finally, to confirm that MICA modifications occur within the NLRP3 PYN domain, we overexpressed the PYN in HEK293T cells treated with CBR-470-2 or MGO. We then excised the HMW band correspond to the PYN dimer and the monomeric PYN domain from SDS-PAGE gels and subjected these isolates to enzymatic digestion. We then monitored the formation of the MICA modification by mass spectrometry. Crosslinked peptides with the corresponding MICA mass addition (36 Da) were observed in isolates prepared from both the CBR-470-2 and MGO treated samples between C08-R43, C08-R81, C38-R07, C38-R43, C108-R89 (CBR only), C130-R89, and C130-R126 and were not observed in the PYN monomer bands isolated from untreated cells (**Fig. 4D-G, Fig. S5A-F**). The formation of crosslinks involving C08 and C38, combined with the minimal increase in HMW bands for PYN constructs containing only these Cys residues (**Fig. 4C**), suggests that these crosslinks are likely intramolecular. This would indicate that apart from the intermolecular crosslinks observed by SDS-PAGE (evident by HMW bands), CBR-470-2 treatment also leads to intramolecular crosslinks. Further, this mix of crosslinks suggests that NLRP3 is highly sensitive to glycolytic perturbations resulting in MGO modification at multiple sites that integrate to inhibit of NLRP3 inflammasome assembly and activity.

## Discussion

Here, we have discovered that inhibition of the glycolytic enzyme PGK1 in a monocyte cell line leads to accumulation of methylglyoxal, a reactive metabolite that covalently crosslinks and inactivates NLRP3, the central pattern recognition receptor of the NLRP3 inflammasome. Interestingly, inhibition of other glycolytic enzymes has been found to modulate the activity of NLRP3 inflammasome, as hexokinase inhibition leads to increases in cytosolic mtDNA which induces inflammasome activation^32,33^. Despite this, regulation of the NLRP3 inflammasome by glycolytic metabolites is poorly understood, as inhibition at different nodes of glycolysis can lead to differing effects on inflammasome activity. For example, inhibiting glycolytic commitment using 2-deoxyglucose can either activate or inhibit the NLRP3 inflammasome^33-35^. Nevertheless, these data along with the work presented here present a mounting case that glycolytic metabolism is intimately linked to NLRP3 inflammasome pathway activity. It is conceivable that certain cell types, like pro-inflammatory macrophages (M2 macrophages), which rely on augmented Warburg-like levels of glycolysis, may sense and respond to modulated glycolytic flux, and the mechanism delineated here may provide a means of feeding back to resolve inflammatory activation.

Inhibition of the NLRP3 inflammasome by increased MGO levels occurs as the result of covalent crosslinking of cysteines and arginines through MICA modification, which leads to formation of NLRP3 oligomers. These crosslinks occur through a variety of cysteines and arginines across multiple domains of NLRP3, having established in this work 7 different crosslinks within the pyrin domain alone. Previously, we reported that VLX1570, a covalent small molecule inhibitor of NLRP3 can also crosslink NLRP3 through its multiple covalent reactive groups. The pyrin domain was also the most sensitive to crosslinking by VLX1570, although, like MGO, we observed crosslinking of the other domains of NLRP3. In both cases, the crosslinking appears to be nonspecific for a specific site in NLRP3, but instead seems to form disordered multimers of various sizes which we posit inhibits the appropriate oligomerization and assembly of the NLRP3 inflammasome. Given the propensity of the pyrin domain, and more generally NLRP3, it is possible that covalent crosslinking and inactivation is an evolved mechanism to sense the presence of an array of reactive metabolites.

Here we have shown that MGO acts similarly to the reactive metabolites fumarate and itaconate, capable of both activating NRF2 and inhibiting the NLRP3 inflammasome. This concurrent signaling may suggest an evolved coordinated response to promote cellular survival and dampen inflammation. To the extent that MGO might modulate additional related cellular responses will be of keen interest in future work. Nevertheless, these data suggest that one reactive electrophilic compound is likely capable of modulating both pathways, potentially for therapeutic benefit in disease. Indeed, there are numerous disease states in which NRF2 activation and NLRP3 inhibition are known to be beneficial including neurodegenerative diseases, metabolic diseases, gastrointestinal diseases, and autoimmune disorders. Using a compound such as CBR-470-2, which beneficially regulates both pathways, may provide an additive or synergistic therapeutic effect to improve disease outcome.

## Methods

Methods generally are as previously described^18^ and are here reviewed in brief with any modifications.

### Compounds, Antibodies, and Plasmids

Ultrapure LPS, E. coli 0111:B4 (Invivogen; Cat. No. tlrl-3pelps) was dissolved in ultrapure water and administered at 1 µg/mL. MCC950 (SelleckChem; Cat. No. S7809) was dissolved in water and administered at 10 µM. Nigericin Sodium Salt (Cayman Chem.; Cat. No. 11437-25) was dissolved in ethanol and administered at 10 µM. CBR-470-1 and CBR-470-2 were synthesized in house as previously described^28^. Methylglyoxal (MGO) was obtained from Acros Organics (40 wt% solution in water, Cat. No. 175791000) and freshly diluted in cell culture media prior to administration at the indicated concentrations.

### Cell Culture and ASC Speck Assay

WT THP1 cells, THP1-ASC-GFP Reporter cells, and HEK-Blue™ IL-1β cells were obtained from Invivogen and maintained according to Invivogen protocols. siRNA transfections were performed using Lipofectamine™ RNAiMAX Transfection Reagent (Invitrogen, 13778075) following manufacturer’s protocol with 9 µL of RNAiMAX reagent and 3 µL of 10 µM siRNA in 300 µL of Optimem per 2 million cells in a 6 well dish. Cells with knocked down PGK1 from siRNA experiments were tested 2 days following transfection in respective assays or harvested for gene knockdown confirmation. For ASC-GFP Speck assay, cells were plated in black, clear-bottom 384 well plates at 20,000 cells/well in 50µL of media. Compounds in dose response were transferred using an Agilent Bravo outfitted with a pintool head to transfer 100 nL. Following pre-treatment time (1 to 24 h), 10 µL of media containing Nigericin and Hoechst were added to each well for two hours (final concentration 10 µM Nigericin, 5 µg/mL Hoechst). Cells were imaged on the Cellinsight CX5 HCS Platform and ASC-Specks were quantified using the SpotDetector function of the Cellinsight High Content Analysis Platform. GSH curve shift assays were performed in a similar manner except 10 µL of GSH (final concentration 10 mM) was added 15 min prior to addition of CBR-470 compounds.

### IL-1β Secretion Inhibition Assay

WT THP1 or primary dendritic cells were primed and treated with compound at the indicated concentrations. Following pre-treatment of 2 h with inhibitors, cells were treated with ATP pH 7.4 (final concentration 5 mM). Cells were allowed to secrete IL-1β overnight. The next day, 10,000 HEK-Blue™ IL-1β cells were plated in 30 µL of media in black, clear bottom 384-well plates, and 10 µL of IL-1β conditioned media added to each well. Cells were incubated overnight to produce SEAP. The following day, 30 µL of QUANTI-Blue (Invivogen) was added to each well and incubated at 37 °C for 30 min-24 h (until visibly observable signal) and absorbance at 655 nM was measured.

### Pyroptotic Cell Death Assay

WT THP1 cells were primed with LPS, pretreated with compounds, and activated with Nigericin as described in the ASC-Speck Assay. 10 µM MCC950 was used as a control to inhibit pyroptotic cell death, and cells not treated with Nigericin were used for no pyroptotic cell death (maximal viability). Two hours after addition of Nigericin, 30 µL of CellTiter-Glo (Promega, diluted 1:6 in water), was added to each well and luminescence was measured with an Envision plate reader.

### qPCR

Cells were lysed and RNA purified using a RNeasy Mini Kit (Qiagen, Cat. No. 74106) following the manufacturer’s protocol. 400 µg of RNA was converted to cDNA using the High-Capacity cDNA Reverse Transcription Kit (Advanced Biosystems; cat. 4368814) which was then diluted 1:4 with DNase/ RNAse free water. qPCR reactions were prepared using Power SYBR Green PCR Master Mix (Applied Biosystems; cat. 4367659) and primers (Table S3) obtained from Integrated DNA Technologies. qPCR reactions were performed using the QuantStudio™ 7 Flex with an initial melting period of 95 °C for 10 min and then 40 cycles of 15 s at 95 °C, 1 min at 60 °C.

### Immunoblotting and Immunoprecipitation

To express NLRP3 in HEK293T cells, 10^6^ cells were plated on 6-well plates coated with poly-d-lysine and transfected with 2 µg of DNA per well and 7 µL of Fugene in 100 µL of Optimem. After 48 h, the cells were treated with compound. For experiments done with endogenous NLRP3, WT THP1 cells were primed overnight with 1 µg/mL LPS, 1 ng/mL TNF-α and treated with compound before collection. Each well was harvested in RIPA buffer with protease inhibitor and lysed on ice for >15 min. Insoluble material was separated via centrifugation. Concentration of soluble lysates was measured via absorbance with a nanodrop instrument. Lysate samples loaded typically were 50 µg. For samples with FLAG immunoprecipitation, lysates were normalized to 1 mg/mL and 20 µL of Magnetic FLAG Beads were added to each sample and incubated at 4°C overnight. Beads were immobilized using a magnetic Eppendorf holder and washed 2X with lysis buffer and 1X with DPBS. Beads were either resuspended in loading dye for analysis via Western blot, or in 100 µL of DPBS for click chemistry. Click Chemistry Master Mix was comprised of 6 µl of 1.7 Tris((1-benzyl-1H-1,2,3-triazol-4-yl)methyl)amine (Sigma) in 4:1 tBuOH:DMSO solution, 2 µl of 50 mM CuSO4 (Sigma) in water, 2 µl of 5 mM rhodamine-azide in DMSO and 2 µl of 50 mM tris(2-carboxyethyl)phosphine (TCEP) in water. To each sample, 12 µL of click chemistry master mix was added and incubated for 1 h in the dark at room temperature. Beads were immobilized using a magnetic Eppendorf holder and washed 2X with DPBS. Beads were resuspended in loading dye for analysis via Western blot and boiled for 15 min at 95 °C. SDS-PAGE and Western blotting protocol as previously described^18^.

### Cellular Methylglyoxal Level Assay

WT THP1 cells were treated with CBR-470-2 or MGO for the indicated timeframes and concentrations. Cells were harvested via centrifugation, rinsed twice with DPBS, and then lysed in 100 µL of DPBS per 1 million cells via probe sonication. Insoluble material was separated via centrifugation. 30 µL of clarified lysate, 50 µL of DPBS, and 10 µL of 100 µM 1,8-diaminonaphthalene were added to each well in a clear 96 well plate and incubated for 30 min. Fluorescence (Ex: 355 nm, Em: 400 nm) was measured using a SpectraMax iD3 plate reader.

### Mass Spectrometry

The gel band sample was destained, reduced (10 mM DTT), alkylated (55mM iodoacetamide) and digested with trypsin overnight before being analyzed by nano-LC-MS/MS. The nano LC-MS/MS analysis was carried out on a Thermo Scientific Easy nano LC 1200 coupled with Thermo Scientific Q Exactive Plus using Nanospray Flex ion source. Eight μL of digested peptides were analyzed by reverse-phase chromatography using 14cm length x 75mm inner diameter nanoelectrospray capillary column packed in-house with Phenomenex Aqua 3 µm C18 125 Å. Mobile phase A and B were water + 0.1% formic acid and 80% acetonitrile + 0.1% formic acid. The elution gradient started at 2% B for 5min, ramped up to 25% B for 100mins, 40%B for 20 mins, 95%B for 1 min and held at 95% B for 14 mins. Data acquisition was performed using Xcalibur (version 4.3). One MS scan of m/z 400-2000 was followed by 10 MS/MS scans on the most abundant ions with application of the dynamic exclusion of 10sec.

### Recombinant PYN Domain Expression

As previously described^18^

**Table S1:** Compounds

| Compound | Source | Catalog Number |
| --- | --- | --- |
| ML385 | Cayman Chemical | 21114-5 |

**Table S2:** Antibodies for Western Blotting

| Protein | Supplier | Catalog Number | Host Species | Dilution |
| --- | --- | --- | --- | --- |
| NLRP3 (NBD) | Cell Signaling | 15101S | Rabbit | 1:1000 (BSA) |
| NLRP3 (PYN) | Adipogen | AG-20B-0014-C100 | Mouse | 1:1000 (BSA) |
| Tubulin | Sigma | T6557 | Mouse | 1:2000 (BSA) |
| HIS-Tag | Santa Cruz | Sc-8036 | Mouse | 1:1000 (BSA) |
| FLAG | Sigma | F1804 | Mouse | 1:1000 (Milk) |
| IL-1β | GeneTex | GTX74034 | Rabbit | 1:1000 (BSA) |
| GFP | Abcam | Ab290 | Rabbit | 1:1000 (BSA) |
| ASC | Cell Signaling | 13833S | Rabbit | 1:1000 (BSA) |
| CASP1 | Cell Signaling | 3866S | Rabbit | 1:1000 (BSA) |
| NEK7 | Cell Signaling | 3057S | Rabbit | 1:1000 (BSA) |

**Table S3:**
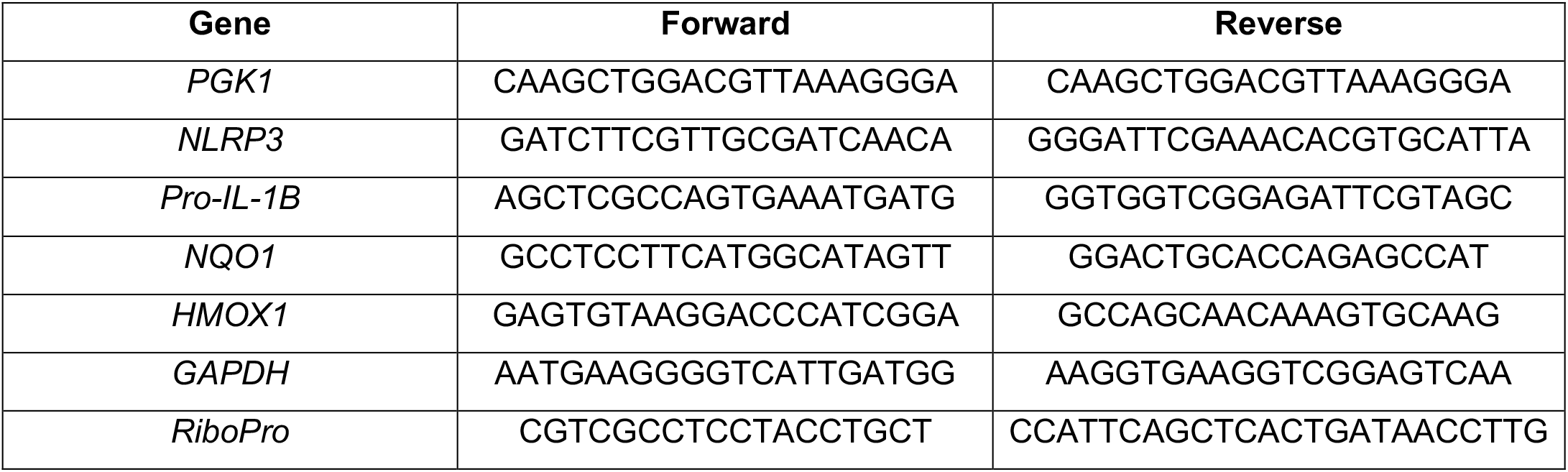
qPCR Primers

## Author Contributions

C.S., R.L.W., and M.J.B. designed research; C.S., C.B., J.S., I.L., and P.B. performed research; C.S., C.B., and I.L. analyzed data; and C.S., R.L.W., and M.J.B. wrote the paper.

## Acknowledgments

This work was supported by the NIH (GM146865 to MJB; DK107604 and AG046495 to RLW). We thank Alan Chu (CALIBR) for providing THP1-ASC-GFP, WT THP-1, and HEK-Blue™ IL-1β cells, Calibr at Scripps Research for providing CBR-470-1 and CBR-470-2, Luke Lairson and lab members for assistance with CX5 HCS platform, and Linh Truc Hoang and the Scripps Mass Spectrometry core for assistance with running and processing mass spectrometry samples,

---

**Supplemental Figure 1.**
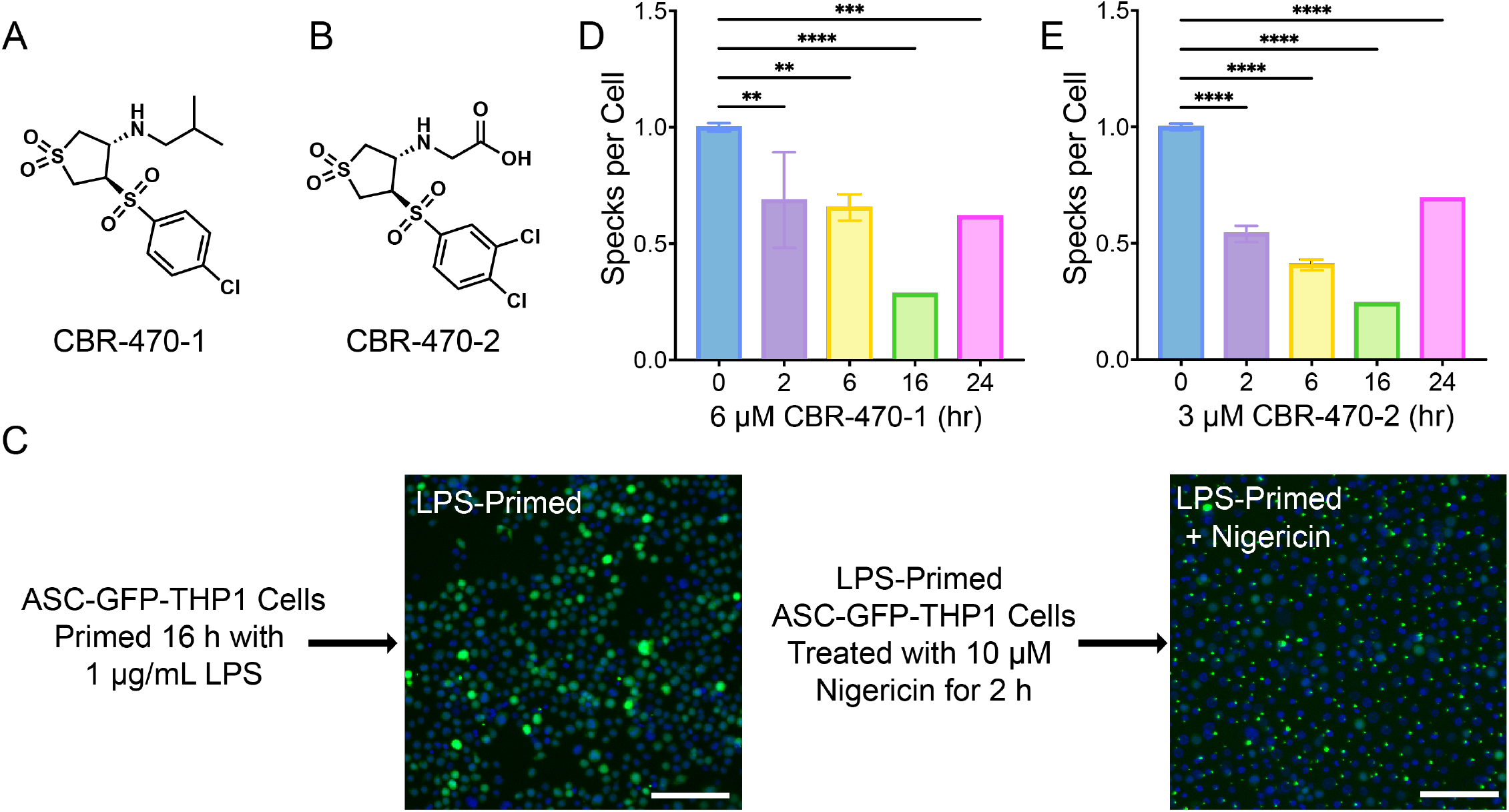
CBR-470-1 and CBR-470-2 display time dependent inhibition at lower concentrations. Structures of CBR-470-1 (A) and CBR-470-2 (B). (C) Schematic with representative images of ASC-GFP Speck formation induced by LPS and Nigericin. Scale bar = 100 µM. (D,E) Number of ASC-GFP specks per cell in THP1-ASC-GFP cells pretreated with CBR-470-1 at 6 µM (D) and CBR-470-2 at 3 µM (E) for 0-24 h. Error bars show SEM for n = 3 replicates.

**Supplemental Figure 2.**
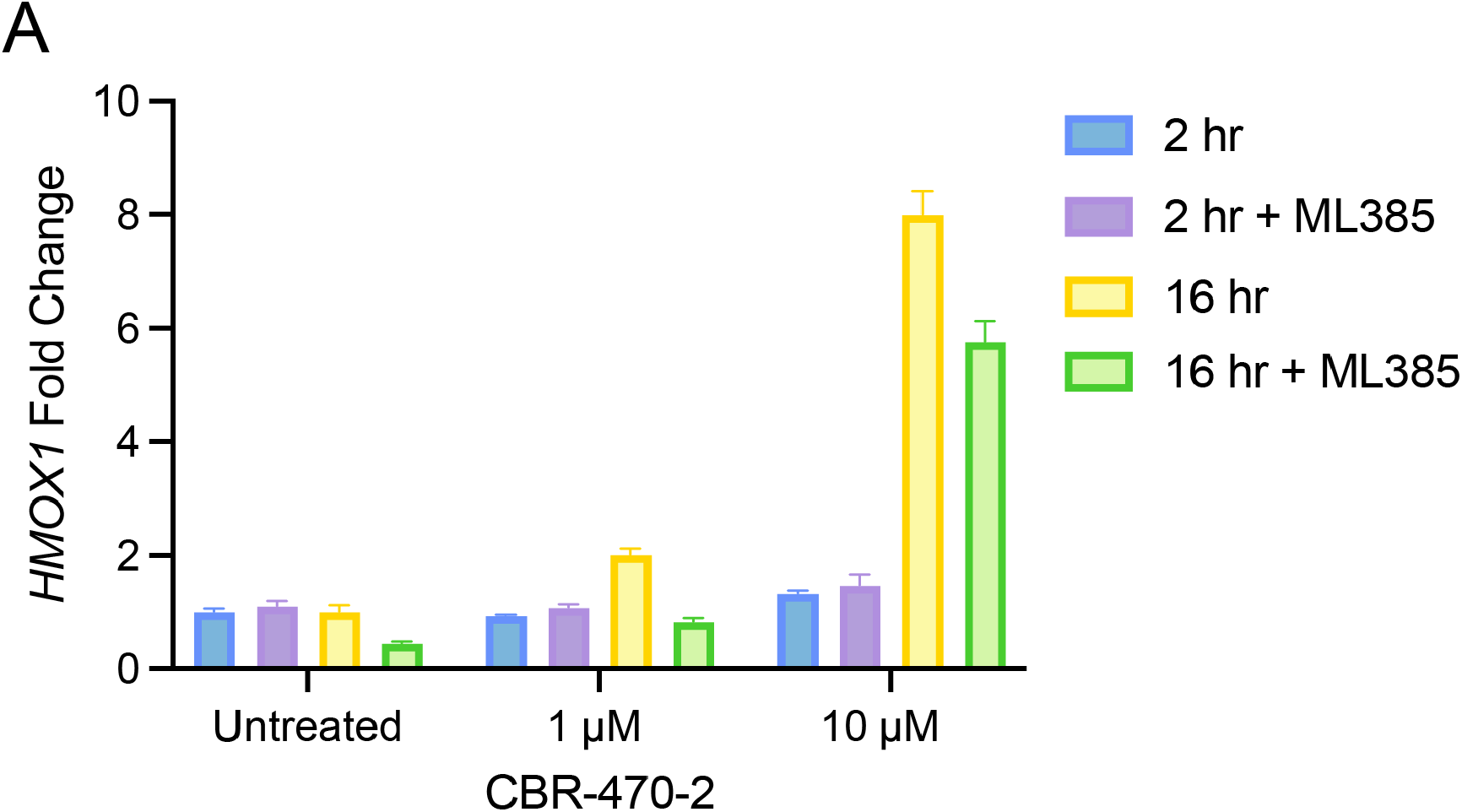
CBR-470-2 induces NRF2 target gene expression at 16 h but not 2 h. (A) Relative transcript level for *HMOX1* as measured by qPCR from WT THP1 cells pre-treated with or without 10 µM ML385 for 30min and then treated CBR-470-2 for 2 or 16 h. Error bars show SEM for n = 3 replicates.

**Supplemental Figure 3.**
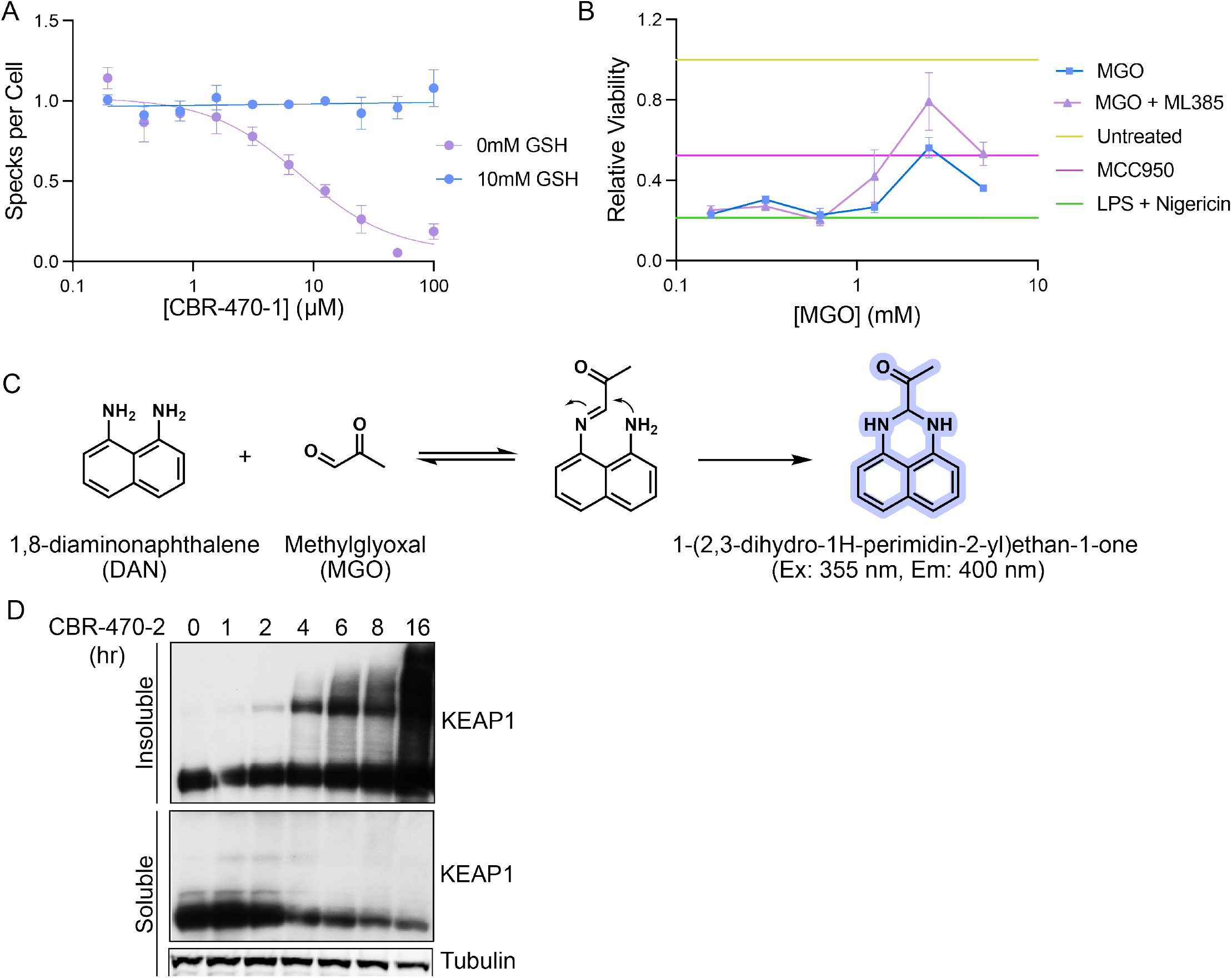
Inhibition or knockdown of PGK1 induces methylglyoxal accumulation. (A) Number of ASC-GFP specks per cell in THP1-ASC-GFP cells pre-treated with 0 or 10 mM GSH and then treated in dose response with CBR-470-1. Error bars show SEM for n = 3 replicates. (B) Relative viability of LPS-primed (1 µg/mL, 16 h) WT THP1 following NLRP3-mediated pyroptotic cell death induced by Nigericin (10 µM, 2.5 h), pretreated with or without 10 µM ML385 for 30 min and then with MGO (2 h) in dose response, or with 10 µM MCC950 (2 h). Error bars show SEM for n = 3 replicates. (C) 1,8-Diaminonapthalene reaction with MGO to form fluorescent compound. (D) Western blot of soluble and insoluble KEAP1 and Tubulin from WT THP1 cells treated with 50µM CBR-470-2 for 1 to 16 h.

**Supplemental Figure 4.**
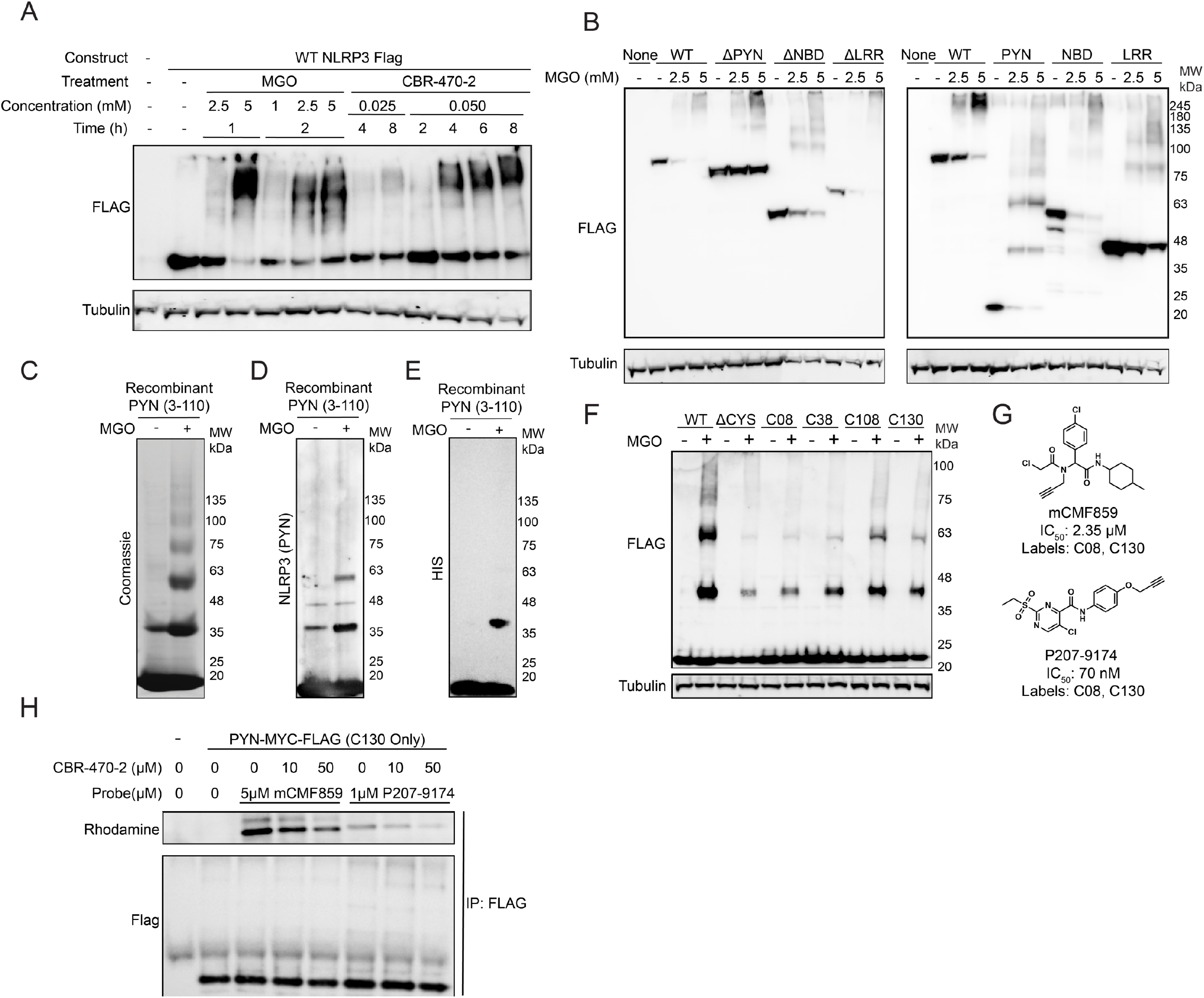
CBR-470-2 and MGO induce covalent crosslinks of NLRP3 Pyrin domain cysteines. (A) Western blot for FLAG and Tubulin in HEK293T cells overexpressing the indicated FLAG-Tagged NLRP3 treated with CBR-470-2 or MGO at the indicated concentrations and timepoints. (B) Western blot for FLAG and Tubulin in HEK293T cells overexpressing the indicated FLAG-Tagged NLRP3 domain constructs treated with 0, 2.5 or 5 mM MGO for 1 h. (C-E) Coomassie Blue stain (C) or Western blots of NLRP3 (PYN) (D) and HIS-TAG (E) from recombinant NLRP3 PYN (3-110, C08S, C38S) treated with 5 mM MGO at 4 °C for 1 h. (F) Western blot of FLAG from HEK293T cells overexpressing NLRP3-FLAG PYN domain constructs with cysteines mutated and individually reintroduced, treated with or without 5 mM MGO for 1 h. (G) Structures and activities of mCMF859 and P207-9174. (H) Anti-FLAG Western blot and rhodamine imaging of FLAG immunoprecipitated material after in situ treatment of HEK293T cells expressing FLAG-tagged PYN domain C130 only construct treated with CBR-470-2 (6h) and then 1 µM P207-9174 or 5 µM mCMF859 (1 h).

**Supplemental Figure 5.**
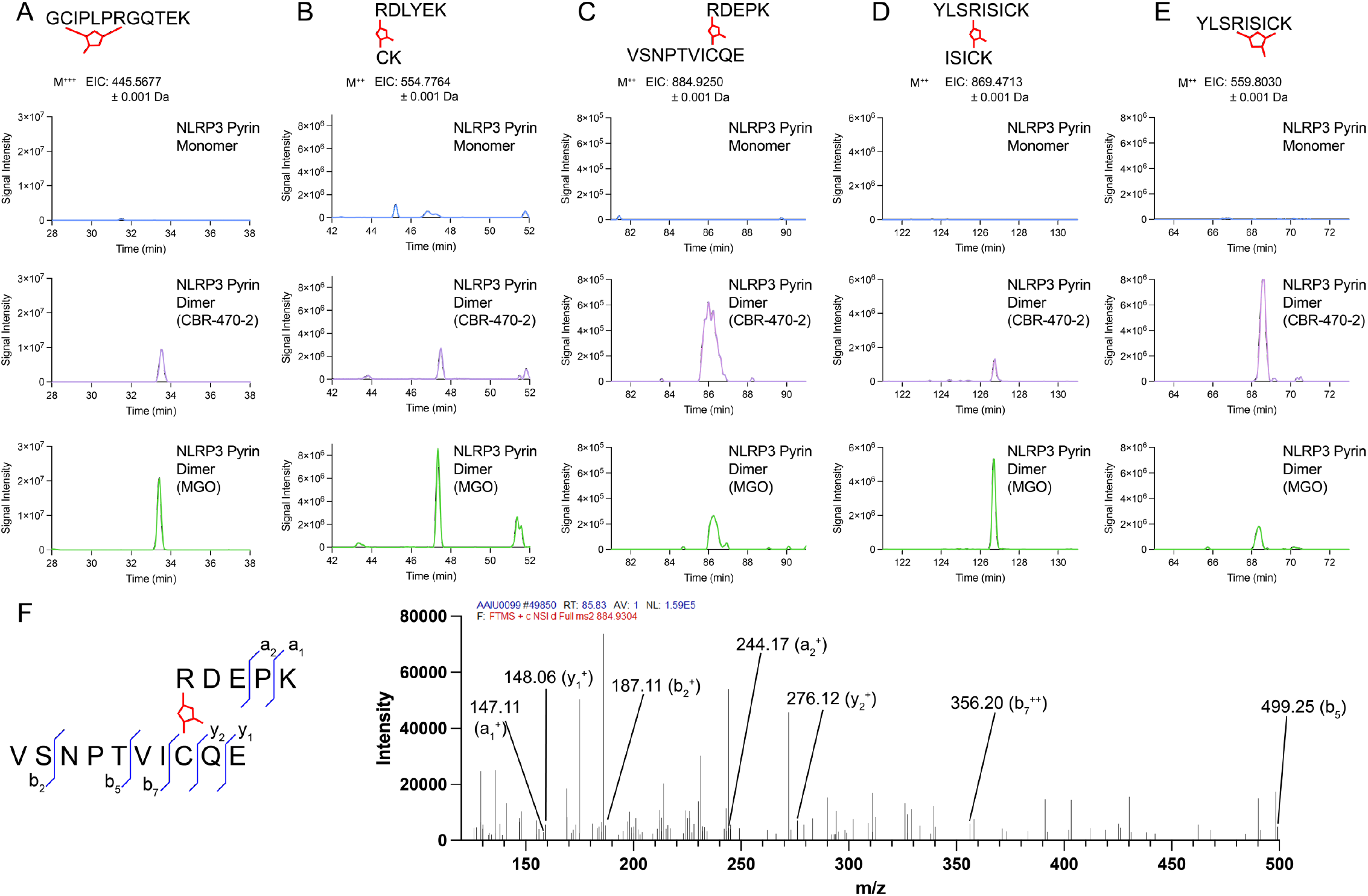
CBR-470-2 and MGO induce MICA crosslinks among NLRP3 Pyrin domains. (A-E) EICs from LC–MS/MS analyses of gel-isolated and digested HMW-PYN (CBR-470-2 and MGO-induced) and monomeric PYN for intramolecular C38-R43 (A), C08-R81 (B), C108-R89 (C), intermolecular C130-R126 (D), and intramolecular C130-R126 (E) crosslinked peptides. (F) Annotated MS2 spectrum from the crosslinked C108-R89 PYN peptide.

